# Integration of high-content fluorescence imaging into the metabolic flux assay reveals insights into mitochondrial properties and functions

**DOI:** 10.1101/758110

**Authors:** Andrew C. Little, Ilya Kovalenko, Laura E. Goo, Hanna S. Hong, Samuel A. Kerk, Joel A. Yates, Vinee Purohit, David B. Lombard, Sofia D. Merajver, Costas A. Lyssiotis

## Abstract

Metabolic flux technology with the Seahorse bioanalyzer has emerged as a standard technique in cellular metabolism studies, allowing for simultaneous kinetic measurements of respiration and glycolysis. Methods to extend the utility and versatility of the metabolic flux assay would undoubtedly have immediate and wide-reaching impacts. Herein, we describe a platform that couples the metabolic flux assay with high-content fluorescence imaging to simultaneously enhance normalization of respiration data with cell number; analyze cell cycle progression; quantify mitochondrial content, fragmentation state, membrane potential, and mitochondrial reactive oxygen species. Integration of fluorescent dyes directly into the metabolic flux assay generates a more complete data set of mitochondrial features in a single assay. Moreover, application of this integrated strategy revealed insights into mitochondrial function following PGC1a and PRC1 inhibition in pancreatic cancer and demonstrated how the Rho-GTPases impact mitochondrial dynamics in breast cancer.

## INTRODUCTION

A primary output of cellular metabolism is chemical energy in the form of adenosine triphosphate (ATP). The major bioenergetic pathways that generate ATP in a cell are glycolysis and mitochondrial respiration^1,2^. Production of ATP from the mitochondria is coupled to the generation of reducing power in the tricarboxylic acid (TCA) cycle, and the respiration-dependent formation of a proton gradient by the electron transport chain (ETC).

While ATP generation is the most well-described role of the mitochondria, this multi-purpose organelle performs several additional important cellular functions^3–6^. Metabolic outputs of mitochondrial metabolism directly regulate signal transduction, gene expression, and cell death. Indeed, it is now established that mitochondrial metabolism impacts numerous physiological and pathophysiological states, including development, cancer, immune recognition and surveillance, and blood glucose control to name a few^6–10^. The widespread influence of mitochondria in health and disease underscores the importance of continued development of strategies to fully characterize mitochondrial metabolism and function.

While recent emerging technologies have permitted more precise examination of mitochondrial functions and properties, each of these techniques are typically performed independently^11,12^. This is problematic in the sense that mitochondria are rapidly undergoing (bio)chemical, morphological, physiochemical, thermodynamic, and other changes at any given time; making experiment-to-experiment comparisons challenging if the measurements are not simultaneous. Therefore, capturing bioenergetic and functional data in a single multifunctional assay has the potential to yield greater, more controlled, and more precise mitochondrial information. Here we describe an integrated platform that utilizes bioenergetic profiling technology alongside imaging of mitochondrial functions and properties to formulate a more complete data set from a single experiment.

Metabolic flux technology using the Seahorse Bioanalyzer has emerged as an industry standard to assess the bioenergetic state of cells *in vitro/ex vivo*. It simultaneously measures pH and oxygen concentration in media as a function of time. These measurements provide a baseline surrogate for glycolytic activity as extracellular acidification rate (ECAR), that is media pH reflecting lactic acid abundance, and mitochondrial respiration or oxygen consumption rate (OCR), as determined by the extracellular oxygen level. In addition, metabolic flux technology can also provide information on the bioenergetic properties and functional status of mitochondria. For example, mitochondrial poisons can be used to infer the bioenergetic flexibility of a cell, activity of ETC complexes, and maximal respiration capacity. Indeed, the simplicity, convenience, robustness, and sensitivity of the metabolic flux assay have made it a technology of choice for many laboratories^13–17^.

Despite the widespread use of the Seahorse Bioanalyzer technology, acquisition of reliable data requires effective normalization strategies to correct for cell density. Multiple normalization methodologies have been used with varying degrees of acceptance by the research community. Examples include normalization to post-assay protein harvest or post-assay cell counting, normalization to pre-assay cell counting^18^, or normalization via one of a variety of chemical colorimetric or fluorometric readouts (e.g. MTT, ATPGlo, WST-1). Recent adaptations to the metabolic flux assay have incorporated nuclei fluorescent staining (Hoechst dye) and subsequent imaging with the BioTek Cytation 5 (BioTek, VT, USA) to more adequately control for cell number^19^. Herein, we optimize and extend this concept, as nuclei staining with DAPI or Hoechst dyes can also be applied to determine cell-cycle analysis^20,21^; an important cellular characteristic when examining drug responses in high-throughput screens. Importantly, mitochondrial bioenergetics have been previously shown to coordinate with cell cycle dynamics^22,23^, further supporting the use of nuclei counterstaining in conjunction with the metabolic flux assay.

The use of nuclear stains can be valuable not only for cell counting, but for generation of nuclei masks to permit localized image analysis of fluorescent events in a distance constrained fashion (e.g. 10μm from nuclei mask); allowing quantification of secondary fluorescent signals at single-cell resolution. For instance, we have applied fluorescent staining of mitochondria via the MitoTracker Red cell dye and use proximity to the nucleus to quantify mitochondrial content. MitoTracker Red is actively sequestered and retained in mitochondria^24^, a process that is initially dependent on an intact mitochondrial membrane potential^25,26^. In addition to basic identification and quantification of mitochondria, we utilize MitoTracker dye in combination with high-content imaging to analyze mitochondrial fragmentation as an added feature integrated into our analysis pipeline^26,27^.

Additional fluorescence-based dyes are similarly available to measure discrete mitochondrial parameters, including membrane potential (ΔΨ_m_) and mitochondrial reactive oxygen species (mtROS). ΔΨ_m_ is generated by the proton pumping complexes of the ETC. The energy “stored” in the ΔΨ_m_ is ultimately used to drive ATP production by complex V. While moderate fluctuations in the ΔΨ_m_ can reflect normal functioning of mitochondria, sustained increases or drops can lead to mitochondrial pathology and/or target mitochondria for degradation. To build in the detection of ΔΨ_m_ into our imaging platform, we utilized the fluorescent dye tetramethylrhodamine ethyl ester (TMRE). TMRE is sequestered in the inner membrane space (IMS) by active mitochondria based on its negative charge. Depletion of the ΔΨ_m_ leads to loss of sequestration and signal.

Mitochondria are one of the primary sources of reactive oxygen species (ROS), which have been characterized to play important roles in physiological and pathophysiological processes^28–30^. The partial reduction of oxygen by the ETC leads to the formation of superoxide, a potent mitochondrial ROS (mtROS). MitoSOX is a mitochondrially targeted fluorescent dye^11^ that is oxidized to ethidium by superoxide. The ethidium then intercalates into mitochondrial DNA and thus produces fluorescence^31^.

In this report, we describe a method that integrates the analysis of mitochondrial bioenergetics with mitochondrial properties, by implementing a variety of chemical fluorescent stains and high-content imaging into the Seahorse metabolic flux assay. This includes image analysis for cell number normalization and cell cycle distribution, along with mitochondria quantity, localization, fragmentation, membrane potential, and ROS (**Fig. 1**). We established the utility of this novel methodology in human breast and pancreatic cancer cell lines using a variety of pharmacological probes, including those that perturb nuclear content and mitochondrial functions, and respiration deficient pancreatic cancer cells. Then, we further extended the utility of our platform by interrogating mitochondrial content and function following genetic knockdown of the mitochondrial master regulatory proteins PGC1a and PRC1. Finally, application of this strategy yielded insights into the role of Rho-GTPases in mitochondrial dynamics in breast cancer. While the roles of the Rho-like family of Rho-GTPases in breast cancer progression were well characterized, our study is the first to determine their role in the regulation of mitochondrial content, fragmentation, and respiratory capacity^32–36^. Collectively this study enhances the utility of the metabolic flux assay and provides a more complete platform to study mitochondrial biology from multiple dimensions, simultaneously.

**Fig. 1.**
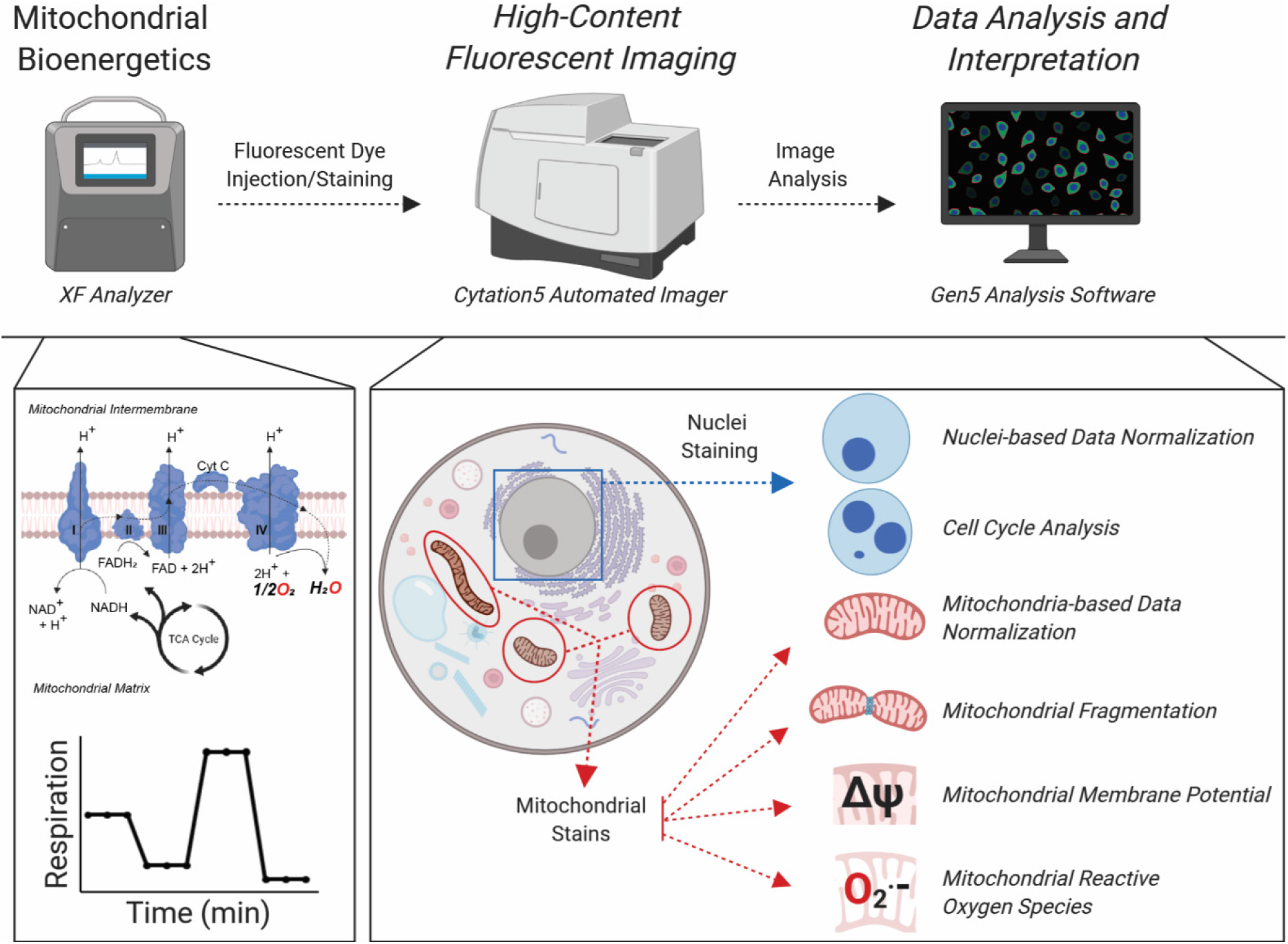
Platform to integrate the metabolic flux assay with high content imaging. Schematic overview of the integration of the metabolic flux assay with high-content fluorescent imaging and data analysis workflow. At the instrument level, cells are processed using the Seahorse metabolic flux assay and immediately stained with a variety of nuclear and mitochondrial dyes, which is completely integrated in the Seahorse bioanalyzer assay. The plates are then abstracted and imaged on a Cytation5 Automated Imager for downstream image analysis and interpretation. At the biochemical level, the metabolic flux assay provides OCR and ECAR data and information on other mitochondrial bioenergetic properties (by Mito Stress Test). The cells are then stained with nuclear and mitochondrial dyes that provide information on the cellular properties noted.

## METHODS

### Cell culture

PA-TU-8902, MIA PaCa2, S2-013, and T3M4 pancreatic cancer cell lines were cultured in DMEM supplemented with 10% FBS. S2-013 ρ^0^ cells were supplied additionally with 10 μg/ml EtBr, 100 μg/ml pyruvate and 50 μg/ml uridine. The VARI068 breast cancer cell line was maintained in RPMI1640 supplied with 10% FBS. Cell lines were STR profiled and routinely tested for mycoplasma.

### Chemicals and probes

FCCP, oligomycin, rotenone, antimycin A, TBH70X, tert-Butyl hydroperoxide solution (Luperox), taxol, and Poly-L-Lysine (mol wt 70,000-150,000, 0.01%) were obtained from Sigma. Hoechst 33342, Mitotracker DeepRed, MitoSOX, and SYTOX Green were from ThermoFisher Scientific. TMRE was from Abcam. All compounds were stored at −20°C except Taxol and Hoechst (4°C). Dyes were stored protected from light. FCCP, Rotenone, Antimycin A, Hoechst, MitoTracker DeepRed, and SYTOX were stored at −20°C, TMRE was pre-diluted at 10μM in media (10X) and aliquoted for single use. MitoSOX was likewise aliquoted for single use.

### shRNA constructs and viral transduction

pLKO lentiviral vectors were ordered as bacterial glycerol stocks from Sigma, MISSION® shRNA Bacterial Glycerol Stock, cat# SHCLNG-NM_013261. The shNT sequence was subcloned into pLKO backbone vector. The references for the sequences are provided in Supplementary Table 1. Viral particles were produced by the University of Michigan Vector Core. T3M4 cells were transduced with the addition of polybrene (Sigma) to 8μM final concentration. Cells were selected with 1 μg/mL puromycin (Sigma) for 3 days. Following selection, transduced T3M4 cells were seeded at 20,000 cells/well for the metabolic flux-imaging assay as described below. Remaining cells were processed for qPCR.

### RNA isolation and qPCR

10^6^ cells were lysed and RNA isolated using the RNAeasy kit (Qiagen) according to the manufacturer’s instructions. 1μg of total RNA was added for cDNA synthesis using the iScript cDNA Synthesis Kit (Bio-Rad) following the manufacturer’s protocols. For qPCR, Fast SYBR Green Master Mix (ThermoFisher Scientific) was used, and amplification was detected with an Applied Biosystems QuantStudio3 Real-Time PCR System. The sequences of the primers used for the amplification are provided in the Supplementary Table 2.

### Metabolic flux assays

Adherent cells were seeded at 2×10^4^ cells/well in normal growth media (cell line specific) in a Seahorse XF96 Cell Culture Microplate. To achieve an even distribution of cells within wells, plates were rocked at 25°C for 20-40 minutes. For each staining group, one extra well on the outer perimeter of the plate was seeded to calibrate image acquisition parameters. The plate was then incubated at 37°C overnight to allow the cells to adhere. The following day, growth media was exchanged with Seahorse Phenol Red-free DMEM and either basal OCR was measured (for wells that were to be imaged with mitochondrial dyes, see Imaging section below) or an XF Cell Mito stress test (Agilent) was performed according the manufacturer’s instructions. In both cases, the last injection port was used for cell stain/dye injection. Upon completion of the Seahorse assay, cells were washed three times with pre-warmed phenol-free DMEM media (no FBS) and transferred to the Cytation5 for imaging.

For cells grown in suspension, 10^5^ cells/well were added to a poly-lysine coated Seahorse XF96 Cell Culture Microplate in Phenol Red-free DMEM based assay media. The plate then was centrifuged at 300g for 30 minutes with gentle acceleration and deceleration. The plate is then rotated 180° and the centrifugation repeated. Immediately following completion of the centrifugation, OCR was measured or an XF Cell Mito stress test was performed followed by imaging in the Cytation5 as above; schematic of assay workflow can be visualized in **Supplemental Figure 1A**.

### High Content Imaging

Imaging was carried out using a Cytation5 Cell Imaging Multi-Mode Reader (BioTek, VT, USA). The environment was controlled at 5% CO_2_ and 37°C. Hoechst (1μg/mL final concentration) was imaged using a 365nm LED in combination with an EX 377/50 EM 447/60 filter cube. SyTOX Green was imaged using a 465nm LED in combination with an EX 469/35 EM 525/39 filter cube. TMRE (1μM final concentration) and MitoSOX Red (5μM final concentration) were imaged using a 523nm LED in combination with an EX 531/40 EM 593/40 filter cube. MitoTracker DeepRed (200nM final concentration) was imaged using a 623nm LED in combination with an EX 628/40 EM 685/40 filter cube. Dyes were delivered at the end of the XF Cell Mito stress test from Port D at 10X to the entire plate. Cytation 5 image excitation/emissions spectra utilized for imaging of various fluorescent stains are depicted in **Supplemental Figure 1B.** Image analysis was completed using Gen5 software (BioTek). Example of Gen5 nuclei masking and secondary fluorescent signal masking algorithms can be observed in **Supplemental Fig. 1C**.

## RESULTS

### Integration of fluorescent-based nuclear imaging with the Seahorse metabolic flux assay

#### Normalization of Cell Number

The Seahorse metabolic flux assay is a rapid and robust methodology to measure oxygen consumption rate (OCR) and extracellular acidification rate (ECAR) of living cells in culture. Due to the sensitivity of the Seahorse XF analyzer to measure small changes in OCR and ECAR, it is critical that data are adjusted to account for well to well variability in cell number. To this end, we set forth to develop a high-content fluorescent imaging-based strategy using nuclear staining to quantify cell number directly after the metabolic flux assay (**Fig. 1**). In this iteration of our platform, we first run the Seahorse Mito Stress Test assay, in which OCR is measured at baseline and then following sequential administration of mitochondrial poisons from the instrument ports. After completion of the assay, we deliver the nuclear staining Hoechst dye via the fourth, and otherwise empty, port to live cells. The plate is then washed, and nuclei are counted on a Cytation5 Cell Imaging Multi-Mode Reader.

Initially, to compare the accuracy of our nuclei identified through image analysis versus cell seeding densities at plating, we seeded serial diluted T3M4 pancreatic cancer cells (from 1,500 to 50,000 cells/well) in XF 96 well plates for Seahorse analysis, nuclei counterstaining, and fluorescent imaging. As expected, we observe increases in raw OCR and ECAR values with increasing cells seeded (**Fig. 2A**, **Supplementary Fig. 2A,B**). When normalizing OCR values according to cells seeded, we observe significant variation (**Fig. 2B**). In contrast, cell normalization using fluorescently labeled nuclei offered more consistent OCR values in the 3,000 to 50,000 cell seeded range (**Fig. 2C**). We do not observe significant variation in normalized ECAR values between the cell counting and nuclei labeling strategies with increasing cell densities (**Supplementary Fig. 2C,D**).

**Fig. 2.**
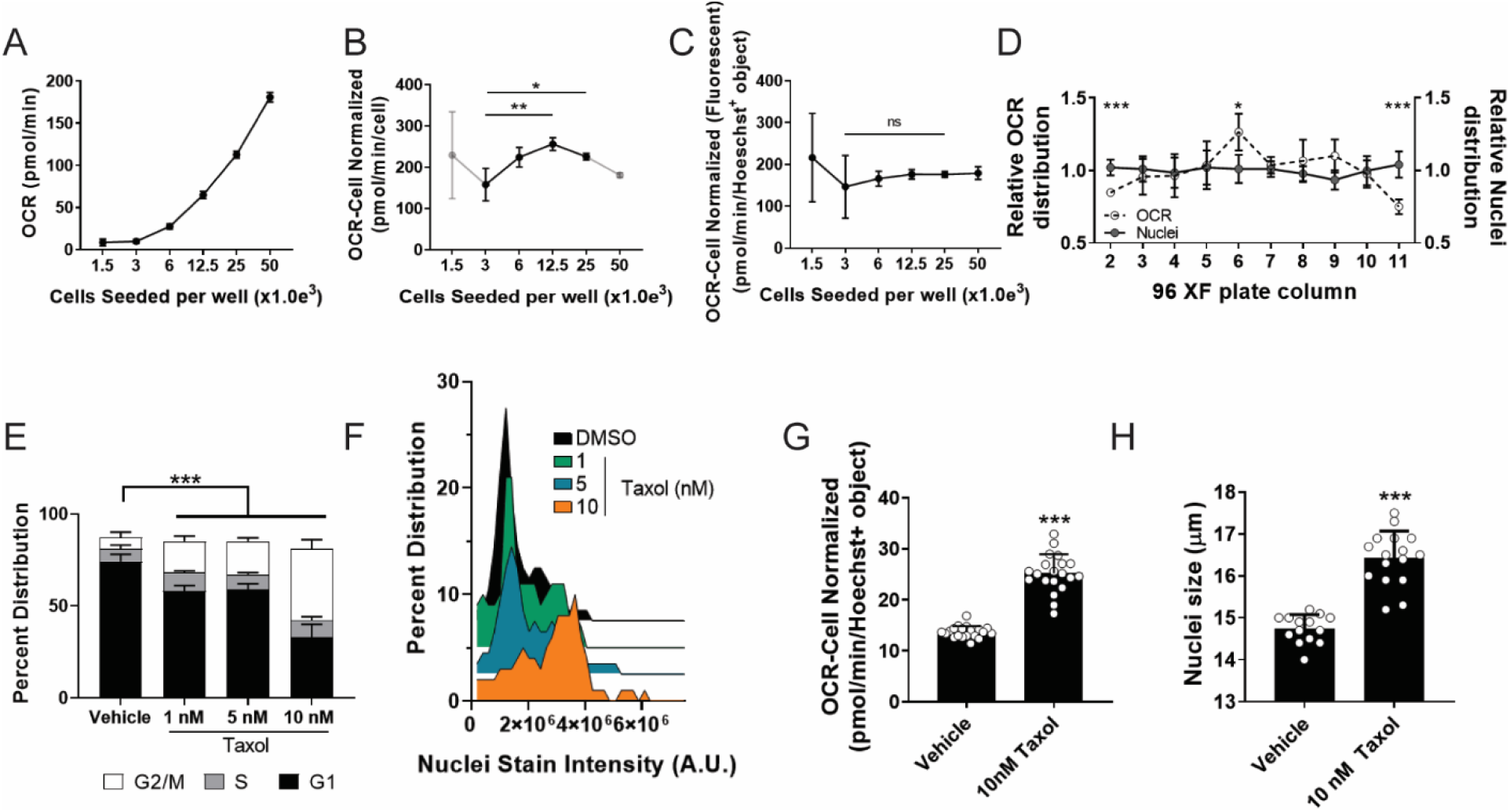
Integration of fluorescent-based nuclear imaging with the Seahorse metabolic flux assay. (**A**) T3M4 cells were seeded in increasing densities in the Seahorse XF plate and processed through the XF assay for OCR and plotted. (**B**) OCR values were corrected for and plotted against manually counted cell seeding densities. Statistical significance determined by one-way ANOVA; *p<0.05, **p<0.01. (**C**) OCR measurements were corrected for fluorescently counted nuclei (i.e. Hoechst+ object) and plotted against seeded cell densities. (**D**) OCR data (empty circle) extrapolated from individual columns of the XF plate display significant variation at the edge columns (i.e. columns 2 and 11), while no changes are observed in nuclei distribution (grey circle). Statistical significance determined by two-way ANOVA; *p<0.05, ***p<0.001. (**E**) Cell cycle analysis distribution of taxol treated cells, identified through nuclei fluorescent imaging (e.g. Hoechst+ object). (**F**) Post hoc distribution analysis of cells treated with increasing concentrations of taxol and their respective nuclei staining intensity (A.U.; arbitrary fluorescent units). (**G**) Image analysis of nuclei size via fluorescently labeled nuclei post taxol treatment. (**H**) OCR values corrected for fluorescently identified nuclei are increased post taxol treatment. Statistical significance in **G,H** determined by Student’s T-test; ***p<0.001. All experiments were the result of ≥ 2 independent experiments.

Furthermore, when plotting OCR values per nuclei relative to the plate column, we noted discrepancies in data acquired from wells at the edge of the XF 96 well plate (**Fig. 2D**). We hypothesized that this resulted from unequal distribution of cells within the well, and in particular, near to the center of the well where the Seahorse microchamber measures oxygen concentration (**Supplementary Fig. 3A**). Indeed, we found that cells in the wells at the perimeter of the plate are more likely to accumulate at the edge of the wells during the centrifugation process, thus artifactually lowering OCR values. Because we were unable to correct for this artifact using nuclei counting, we employ the interior columns/rows of the Seahorse plate for experimentation. Similarly, for assay well for normalization by nuclei counting, we utilize the cells in the center of the well (schematically represented in **Supplementary Fig. 3B**).

#### Cell Cycle Analysis

Nuclei staining intensity can be used to infer the stage of the cell cycle^20,21^. Therefore, we sought to determine if we could integrate this analysis to the XF-Cytation5 platform. To this end, we treated T3M4 pancreatic cancer cells across a dose range of taxol, an FDA approved chemotherapeutic (e.g. Paclitaxel) which arrests cells in G2/M phase of the cell cycle due to its microtubule polymerization inhibiting properties^37,38^. Of note, at the doses examined taxol exhibited cell cycle arrest without toxicity (**Supplemental Fig. 4A,B**). Nuclear staining intensity was then collected and used to demonstrate that taxol induces a dose-dependent accumulation of cells in the G2/M phase of the cell cycle (**Fig. 2E,F**). Similarly, at the highest dose examined, we also observed a significant increase in nuclear size (**Fig. 2G**), demonstrating another utility for our imaging platform. Finally, we observed increased OCR in T3M4 cells that have been treated with taxol, a previously described feature of taxol administration^20^ (**Fig. 2H**). These data support the utility of nuclei counterstaining and imaging to assess cell cycle and nuclear size post metabolic flux assay.

### Integration of fluorescent high-content imaging of mitochondria into the metabolic flux assay workflow

OCR data provide a readout for total respiration of the cells within a well. This is impacted by both the number and activity of the mitochondria in the cells. The latter is related in part to the structural properties of the mitochondria, where fused mitochondrial networks tend to be more respiratory than fissed mitochondria^39,40^. The location of mitochondria within a cell has also been reported to impact their function^41,42^. Furthermore, heterogeneity in mitochondrial density, structure, location, and function also exists on a cell to cell basis within a population of cells in a well. Therefore, we hypothesized that a more complete understanding of this information would provide considerable utility in accurately characterizing mitochondrial function and OCR output from the metabolic flux assay. We therefore adapted our high-throughput imaging platform to capture these parameters by incorporating a series of mitochondrial dyes, delivered in a manner akin to the delivery of the nuclear dye described above, followed by image analysis.

#### Normalization of OCR Data to Mitochondrial Content

We set forth to image mitochondrial content and localization and to evaluate the relationship between these parameters and OCR. To this end, we empirically tested a panel of Mitotracker dyes with T3M4 pancreatic cancer cells plated in the Seahorse 96XF cell plate. Mitotracker Deep Red was identified as the most consistent and robust probe to visualize mitochondria, as it readily retained bright fluorescence in live cell formats as well as post fixation (**Supplementary Fig. 5**). Specifically, it exhibited the highest signal/background ratio in the Seahorse XF cell plate, was fixable, and readily integrates into our Seahorse workflow.

To demonstrate the utility of this approach, we generated a Rho0 (ρ0) S2-013 pancreatic cancer cell line (S2-013-ρ0) with severely depleted mitochondrial DNA resulting from prolonged culture in ethidium bromide^43,44^. Depletion of mitochondrial DNA results in the loss of a functional ETC and thus the ability of mitochondria to maintain their ΔΨ_m_ or respire. As expected, S2-013-ρ0 cells completely lack a respiratory profile (**Fig. 3A**) and have greatly diminished Mitotracker Deep Red staining (**Fig. 3B; Supplementary Fig. 6A**), the latter of which requires ΔΨ_m_ to be retained in the IMS. No changes in ECAR were detected in S2-013 WT vs ρ0 cells (**Supplementary Fig. 6B**). These results, using an artificial system of mitochondrial depletion, illustrate the utility our imaging platform to capture information on mitochondrial content as well as function.

**Fig. 3.**
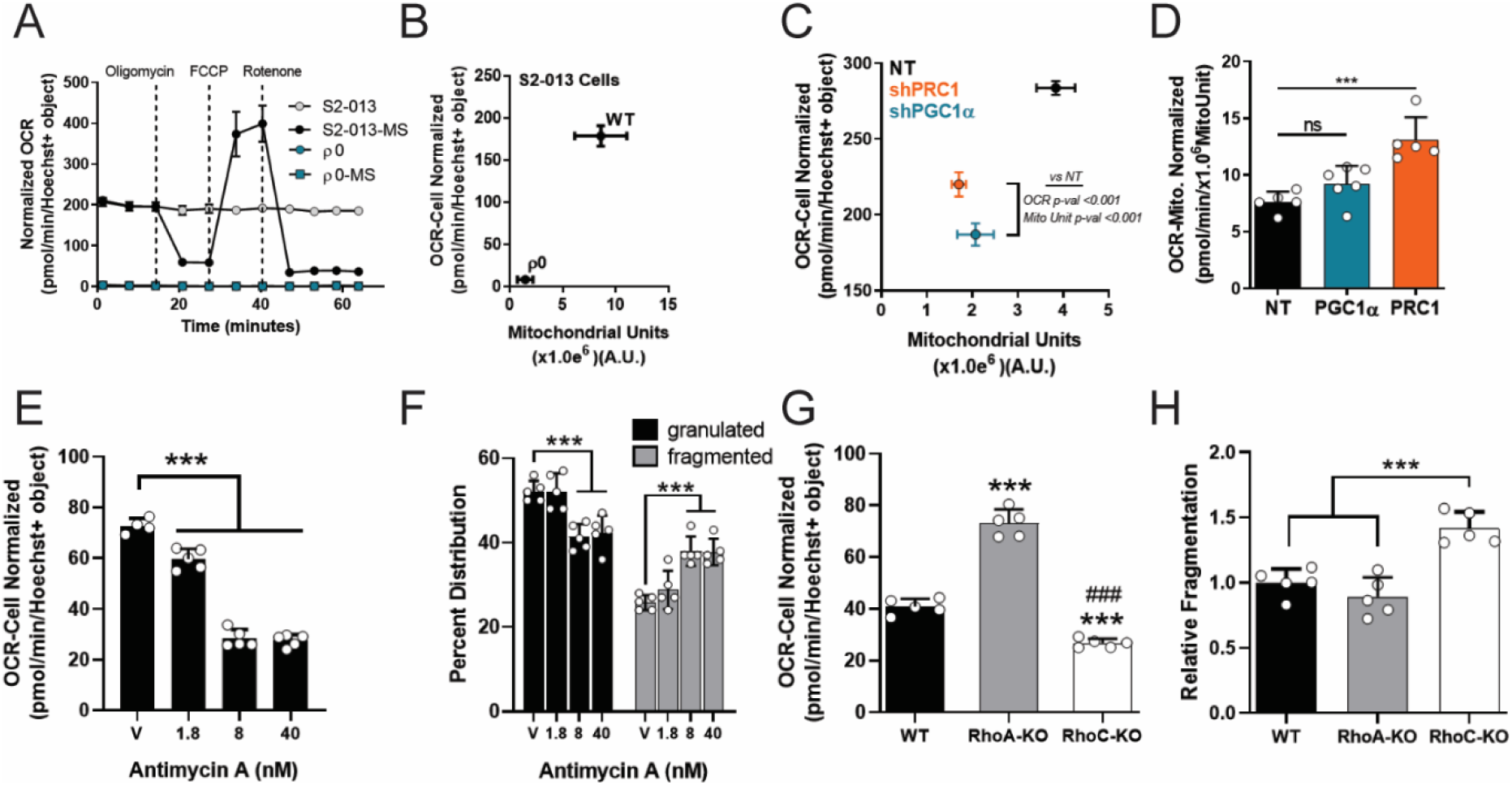
Analysis of mitochondrial quantity and fragmentation downstream of the metabolic flux assay. (**A**) Hoechst+ nuclei corrected OCR data post mitochondrial stress test (“-MS” designation; e.g. S2-013-MS) or measurements of basal OCR values (i.e. S2-013) in wild type S2-013 cells or S2-013-ρ0 (Rho0) cells. (**B**) OCR values corrected for fluorescent nuclei plotted against total fluorescently identified mitochondria (i.e. MitoTracker positive arbitrary fluorescent units; A.U.). (**C**) OCR values normalized for nuclei in shNT (non-targeting), shPRC1, or shPGC1α T3M4 cells, plotted against fluorescently identified mitochondria (MitoTracker positive A.U.). (**D**) OCR values in shNT, shPRC1, or shPGC1α T3M4 cells corrected for total fluorescently identified mitochondria (e.g. Mito Normalized; MitoUnit). Statistical significance determined by one-way ANOVA; n.s., non-significant; ***p<0.001. (**E**) Hoechst+ nuclei corrected OCR values post Antimycin A treatment. Statistical significance determined by one-way ANOVA; ***p<0.001. (**F**) Overall levels of granulated (black) or fragmented (grey) mitochondria post Antimycin A treatment. Statistical significance determined by two-way ANOVA; ***p<0.001. (**G**) Hoechst+ nuclei corrected OCR values of WT, RhoA KO, or RhoC KO VARI068 breast cancer cells. Statistical significance determined by one-way ANOVA; ***p<0.001 vs WT; ###p<0.001 vs RhoA-KO. (**H**) Levels of fragmented mitochondria in WT or RhoA/C-KO VARI068 cells. Statistical significance determined by one-way ANOVA; ***p<0.001. All experiments were the result of ≥ 2 independent experiments.

To extend our imaging platform to a system with a less extreme mitochondrial defect, we employed a genetic approach to deplete mitochondria in a confined timeframe. Protein regulator of cytokinesis 1 (PRC1) and peroxisome proliferator-activated receptor gamma coactivator 1-alpha (PGC-1α) are master regulators of mitochondrial biogenesis^45–48^. We generated PRC1 and PGC-1α knockdown T3M4 cells using shRNA (see qPCR results of gene knockdown in **Supplementary Fig. 7A,B**). Loss of PRC1 or PGC-1α resulted in fewer overall mitochondria, as determined by Mitotracker fluorescent staining and quantification (**Supplementary Fig. 7C**). We then assayed PRC1 and PGC-1α KO cells for basal levels of OCR and normalized the data to nuclei number, followed the XF assay with Mitotracker staining and quantification. Using the nuclei normalization strategy, we observed significantly lower overall OCR values in PRC1 and PGC-1α KO cells as compared to their WT counterparts (**Fig. 3C**). In addition, we find fewer overall mitochondria in the PRC1 and PGC-1α KO cells, as expected (**Fig. 3C**). Surprisingly, however, when we apply normalization of the OCR data at per-mitochondrial resolution (e.g. Mito Normalized; MitoUnit), the normalized data reflects no changes in overall mitochondrial function in PGC-1α KO cells and increased OCR in PRC1 KO cells (**Fig. 3D**). These results suggest that PRC1 and PGC-1α KO cells harbor normally functioning, albeit fewer, mitochondria. This could be easily misinterpreted using traditional normalization strategies. These data strongly support the utilization of mitochondrial quantification as a parallel normalization technique for OCR data in the metabolic flux assay, as the two normalization parameters provide different and important outputs.

#### Mitochondrial Fragmentation Analysis

Next, we sought to explore the use of Mitotracker staining as a method to evaluate mitochondrial fragmentation patterns. Mitochondrial fragmentation has been observed in many settings, including apoptosis, responses to oxidative stress, neurodegeneration, among various others, and is an important feature of mitochondrial biology^49–52^. As a positive control, we treated MIA PaCa-2 pancreatic cancer cells with the ETC complex III inhibitor Antimycin A to induce fragmentation, a previously described feature of its administration^53^. As expected, we observed a decrease in OCR levels in a dose-dependent fashion post Antimycin A treatment (**Fig. 3E**). Using Mitotracker staining/imaging and the BioTek Gen5 spot analysis tool (see BioTek application notes for further detail), we were able to quantify decreases in granulated mitochondria and increases in fragmented mitochondria following Antimycin A treatment (**Fig. 3F**). These results confirmed the utility of mitochondrial image analysis using Mitotracker fluorescent dye for this purpose.

The Rho-like family of Rho-GTPases have been well characterized for their participation in the progression of breast cancer^32,34–36^ and the alteration of metabolic phenotype ^33^. Therefore, we next applied our imaging and analysis strategy to explore the potential for fragmented mitochondria in breast cancer cells that lack either the Rho-like family of Rho-GTPases RhoA or RhoC, via CRISPR/Cas9 knockout (WT, RhoA KO, RhoC KO) (**Supplementary Fig. 8A**). First, we examined basal OCR values in the breast cancer patient-derived cell model VARI068 ^54^ and followed with Mitotracker staining directly into the XF assay. Our data revealed elevated basal OCR levels in the RhoA KO cell line and lower OCR levels in the RhoC KO cell model (**Fig. 3G**). No changes in mitochondrial fragmentation/elongation were observed in the more respiratory RhoA KO cells; in contrast, significantly greater levels of fragmented mitochondria were observed in the RhoC KO cells (**Fig. 3H**). We confirmed that changes in fragmentation were not due to altered overall levels of mitochondria (**Supplementary Fig. 8B**). These data illustrate that quantification of mitochondrial fragmentation via Mitotracker imaging is a robust and useful method that can be readily integrated into the metabolic flux assay to enrich the overall data set regarding mitochondrial features.

#### Fluorescence Staining for Mitochondrial Membrane Potential

To build detection of ΔΨ_m_ into our imaging platform, we utilized the fluorescent dye TMRE. First, we demonstrated that TMRE dye fluorescence correlated with increased OCR values, indicative of active mitochondria (**Fig. 4A**). Uncouplers of ΔΨ_m_, such as FCCP, dissipate the proton gradient. At low concentrations of FCCP, the ETC competes to maintain a proton gradient and induce maximal respiration. At higher concentrations of FCCP, dissipation of the proton gradient outpaces the capacity of the ETC to maintain a gradient, poisoning the mitochondria, and thereby impairing respiration and ΔΨ_m_. Treatment of T3M4 cells with FCCP in a dose-dependent fashion demonstrated this expected rise and fall in OCR (**Fig. 4B**, **Supplementary Fig. 9A**). Similarly, loss of OCR correlated with a loss of TMRE uptake by mitochondria, as reflected by decreased TMRE intensity quantified from fluorescent imagine analysis (**Fig. 4C**); suggesting impaired mitochondrial function.

**Fig. 4.**
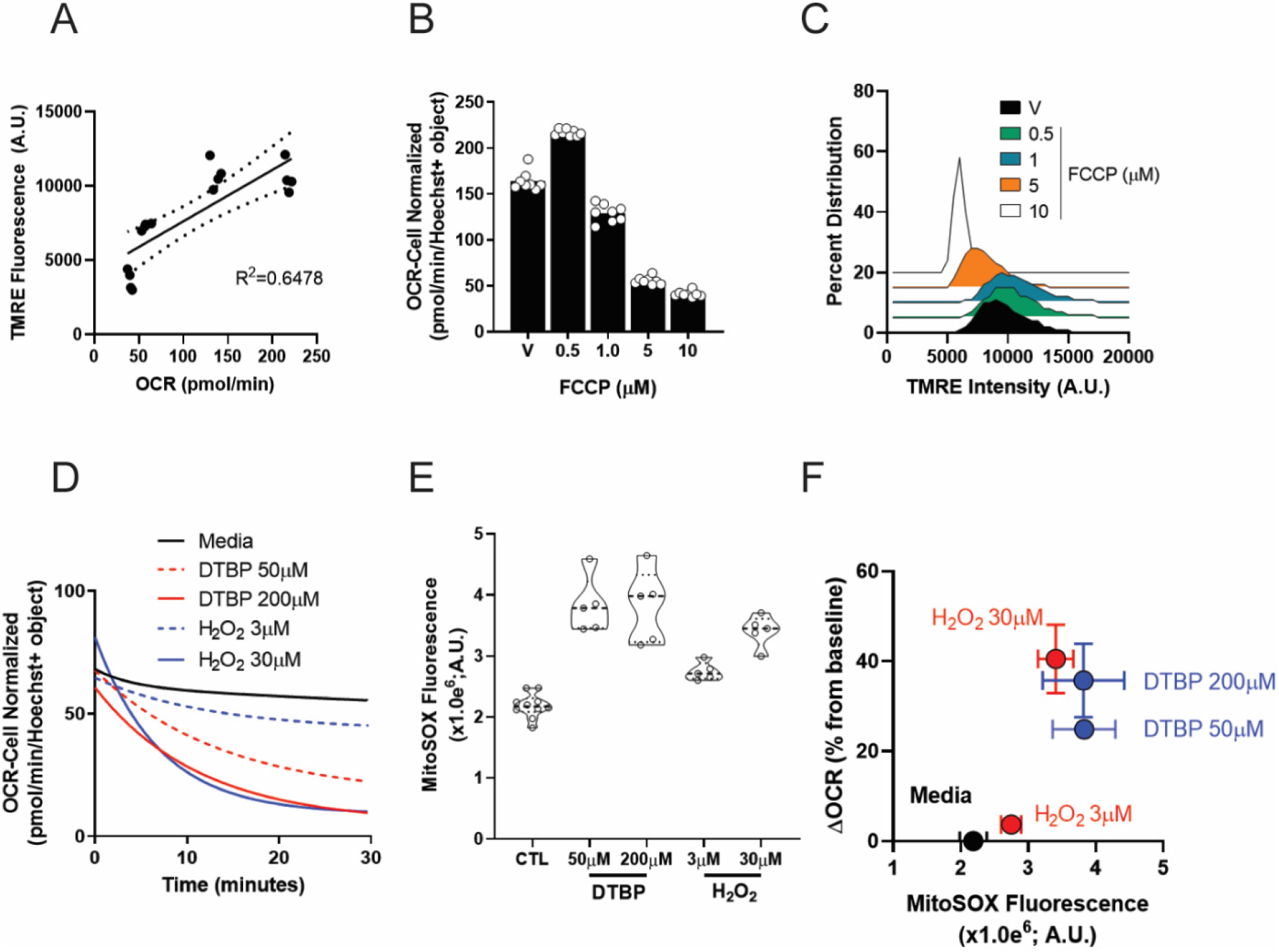
Analysis and quantification of ΔΨ_m_ and mitochondrial ROS. (**A**) TMRE fluorescence plotted with respect to OCR in T3M4 cells. (**B**) OCR readout following treatment across a dose range of FCCP in T3M4 cells. (**C**) Distribution analysis of TMRE fluorescence intensity (A.U.) in T3M4 cells. (**D**) Exponential curve fit of OCR data of PA-TU-8902 pancreatic cancer cells treated with either DTBP (ditertbutyl peroxide; red lines) or hydrogen peroxide (H_2_O_2_; blue lines). (**E**) Violin plots displaying induction of MitoSOX fluorescence (A.U.) following either DTBP or H_2_O_2_ treatment. (**F**) Multi-analysis plot displaying change in OCR values plotted against MitoSOX fluorescence in PA-TU-8902 cells post DTBP or H_2_O_2_ treatment. All experiments were the result of ≥ 2 independent experiments.

#### Mitochondrial Reactive Oxygen Species Fluorescence Imaging

Under physiological conditions, the biogenesis and quenching of ROS regulate various cellular processes and are tightly controlled^55^. Excessive ROS production, on the other hand, is implicated in numerous pathologies, including aging and cancer. Superoxide is the ROS produced from the mitochondria by incomplete oxidation of oxygen during respiration. Therefore, we sought to include detection of mitochondrial superoxide alongside the metabolic flux assay. To measure mitochondrial superoxide, we used the mitochondrially targeted MitoSOX dye. To induce mitochondrial superoxide production, we treated the pancreatic cancer cell line PA-TU-8902 with the membrane-targeted radical initiator ditertbutyl peroxide (DTBP) or hydrogen peroxide (H_2_O_2_). We then monitored OCR levels, integrated MitoSOX dyes into the metabolic flux assay, and observed mitochondrial superoxide via MitoSOX fluorescence. We observed that both H_2_O_2_ and DTBP treatment had a dose-dependent inhibitory effect on OCR levels (**Fig. 4D**). These data are consistent with prior literature demonstrating that oxidative stress can lead to diminished basal OCR values^56,57^. Following DTBP or H_2_O_2_ stimulation, expectedly we quantified a marked increase in MitoSOX fluorescence (**Fig. 4E**) (representative images can be seen in **Supplemental Fig. 9B**). Data corrected for overall changes in OCR versus increases in MitoSOX fluorescence can be observed in **Fig. 4F**. Collectively, these data confirm the utility of MitoSOX dye incorporation into the XF assay, providing robust quantifiable data regarding mitochondrial respiration and the generation of ROS.

## DISCUSSION

Here we present a robust strategy to integrate high-content, high-throughput fluorescent imaging into the Seahorse metabolic flux assay. This integrated approach aims to build upon prior efforts to utilize fluorescence-based counterstains for normalization of bioenergetic data^58^, rather than relying on rudimentary cell number correction strategies. Our pipeline enables the evaluation of many features in one complete experiment on a single XF96 well plate, increasing the utility and output of a single experiment while minimizing the potential for plate-to-plate variability. We initiated this strategy to address observed well to well inconsistencies in the OCR bioenergetic parameter, as similarly reported by others^19,59^. While classic approaches to normalize mitochondrial bioenergetic data with total cellular input or quantified cellular protein may be a quick and efficient, we found these to be inconsistent across experiments and cell lines. Furthermore, as we detail in **Fig. 2**, the location of cells in a well of a Seahorse plate impacts OCR. By accounting for both the number of cells (based on nuclear staining), and more specifically, the number of nuclei in proximity to the sensors, our output provides even greater precision across columns and wells. Furthermore, we employed nuclear size and staining intensity to provide information into cell cycle dynamics.

Extension of OCR normalization to total mitochondria provides further resolution and more granular information about mitochondrial capacity, as illustrated in **Fig. 3**. Staining for total mitochondria is easily achieved through the inclusion of Mitotracker dyes into the metabolic flux assay workflow. We consider this an important experimental control, as a wide array of experimental conditions may impact mitochondrial biogenesis or mitochondrial content, which we have demonstrated affect OCR results. Correction of metabolic flux data using this method provides additional information alongside nuclei-based cell counting. However, the use of Mitotracker dyes in this pipeline are not limited to their use as a normalization tool for metabolic flux data. We also found that the Mitotracker staining patterns readily allow for the identification and quantification of fragmented mitochondria (**Fig. 3E-H**). Indeed, Mitotracker staining could be further amended to observe subcellular mitochondrial localization, organelle or molecular co-localization, fission-fusion dynamics, mitochondrial shape (sphericity), among others.

We then applied this platform in two targeted studies. First, we demonstrated that while knockdown of the master regulators of mitochondrial biogenesis PGC1a and PRC, did result in less cellular respiration, this was the result of fewer mitochondria, not less active mitochondria (**Fig. 3G,H**). This illustrates the utility of building in a mitochondrial normalization strategy into the metabolic flux assay workflow. We then examined how the Rho-like family of Rho-GTPases impact mitochondrial fragmentation, based on their known role in regulating cytoskeletal dynamics. And, more specifically, our previous work found that inhibition of RhoC impairs the metabolic properties of inflammatory breast cancer cells^33^. Application of the MitoTracker strategy downstream of the metabolic flux assay revealed that RhoC deletion significantly increases the number of fragmented mitochondria in inflammatory breast cancer cells, which we hypothesize contributes to the depleted OCR levels (**Fig. 3K,L**). Future studies are required to fully characterize the Rho-like Rho-GTPases and their regulation of mitochondrial dynamics.

The decision to utilize TMRE and MitoSOX dyes were based on the idea of synthesizing a broad data set of mitochondrial features that can be analyzed downstream of mitochondrial respiration on the same cells in the same well. A limitation of the TMRE stain is the requirement for live cell imaging. Subsequent to this, the cells can be stained with MitoTracker, MitoSOX and Hoechst, fixed, and entered into the downstream normalization and analysis workflow. Indeed, we selected a series of dyes with non-overlapping wavelengths (**Supplementary Fig. 1**) so that this suite of parameters can be simultaneously analyzed on a limited number of wells. As noted, MitoTracker, MitoSOX and nuclear staining with Hoechst are amenable to fixation and retain their subcellular localization, and we have had success storing plates for analysis months later. Accordingly, Seahorse plates can be imaged immediately after analysis (post dye addition) or stored for downstream analysis if large numbers of plates are being prepared for high-throughput screening. It is also important to note that cells imaged for MitoTracker, TMRE, and MitoSOX cannot be the same cells on which the Mito Stress Test is performed. Requisite for the Mito Stress Test analysis is the use mitochondrial poisons that impact respiration, membrane potential, oxidative stress, among others. Thus, the staining and analysis protocols need to be performed in parallel.

Finally, we also anticipate that the use of dyes to monitor cellular events beyond those described herein could be readily amended to this workflow. For example, dyes are available to analyze lysosomal labeling (LysoTracker), mitochondrial calcium signaling (e.g. Fura-2, Fluo-3), and/or other oxidative stress parameters (e.g. CellROX)^60^. Similarly, genetically encoded sensors exist to monitor changes in pH, ATP, redox cofactors (e.g. NADH, NADPH), and oxidative stress^61–65^, and these could be applied to this workflow. Indeed, the Cytation5 imaging system permits the use of various filter cube sets allowing visualization of many different fluorescent channels. Finally, this platform also has the potential to be used for drug screening, as approaches to disrupt mitochondrial metabolism for cancer therapy, to facilitate mitochondrial function in aging and other mitochondrial disease, or to promote the turnover of damaged mitochondria in degenerative conditions remain focal points in the development on novel therapeutics^66–69^.

In total, our platform provides high-resolution normalization strategies for Seahorse data that encompass nuclei and mitochondrial based fluorescent imaging and quantification. We define the inclusion of additional mitochondrial stains to generate a robust data set, in one high-throughput experiment, characterizing mitochondrial biology in a continuous kinetic fashion. This is imperative in studying mitochondria, as they can rapidly change phenotypes in response to their environment. Thus, capturing the greatest amount of data in a continuous, rapid manner will provide more consistent results and may reveal additional mitochondrial characteristics otherwise not captured in single experiment formats.

## ACKNOWLEDGMENTS

We would like to thank Dr. John Dishinger of BioTek for his assistance with designing imaging and image analysis protocols. Figure 1 and Supplementary Figure 1A and 3B were created using BioRender.com. C.A.L. was supported by a Pancreatic Cancer Action Network/AACR Pathway to Leadership award (13-70-25-LYSS); Junior Scholar Award from The V Foundation for Cancer Research (V2016-009); Kimmel Scholar Award from the Sidney Kimmel Foundation for Cancer Research (SKF-16-005); a 2017 AACR NextGen Grant for Transformative Cancer Research (17-20-01-LYSS); and an ACS Research Scholar Grant (RSG-18-186-01). D.L. was supported by R01GM101171. A.C.L. and S.D.M. were supported by the Breast Cancer Research Foundation. C.A.L. and S.D.M. were supported by the Rogel Cancer Center core grant NIH-P30-CA046592-29.

## AUTHOR CONTRIBUTIONS

Conceptualization: A.C.L., I.K., C.A.L.; Methodology: A.C.L., I.K., J.A.Y.; Investigation: A.C.L., I.K., L.E.G., H.S.H., S.A.K., J.A.Y., V.P.; Visualization: A.C.L., I.K.; Formal Analysis: A.C.L., I.K.; Manuscript and Figure Preparation: A.C.L, I.K., C.A.L.; Resources: D.L., S.D.M., C.A.L.; Supervision: S.D.M., C.A.L.; Funding Acquisition: S.D.M., C.A.L.

