## Supplemental Information for "Integration of high-content fluorescence imaging into the metabolic flux assay reveals insights into mitochondrial properties and functions"

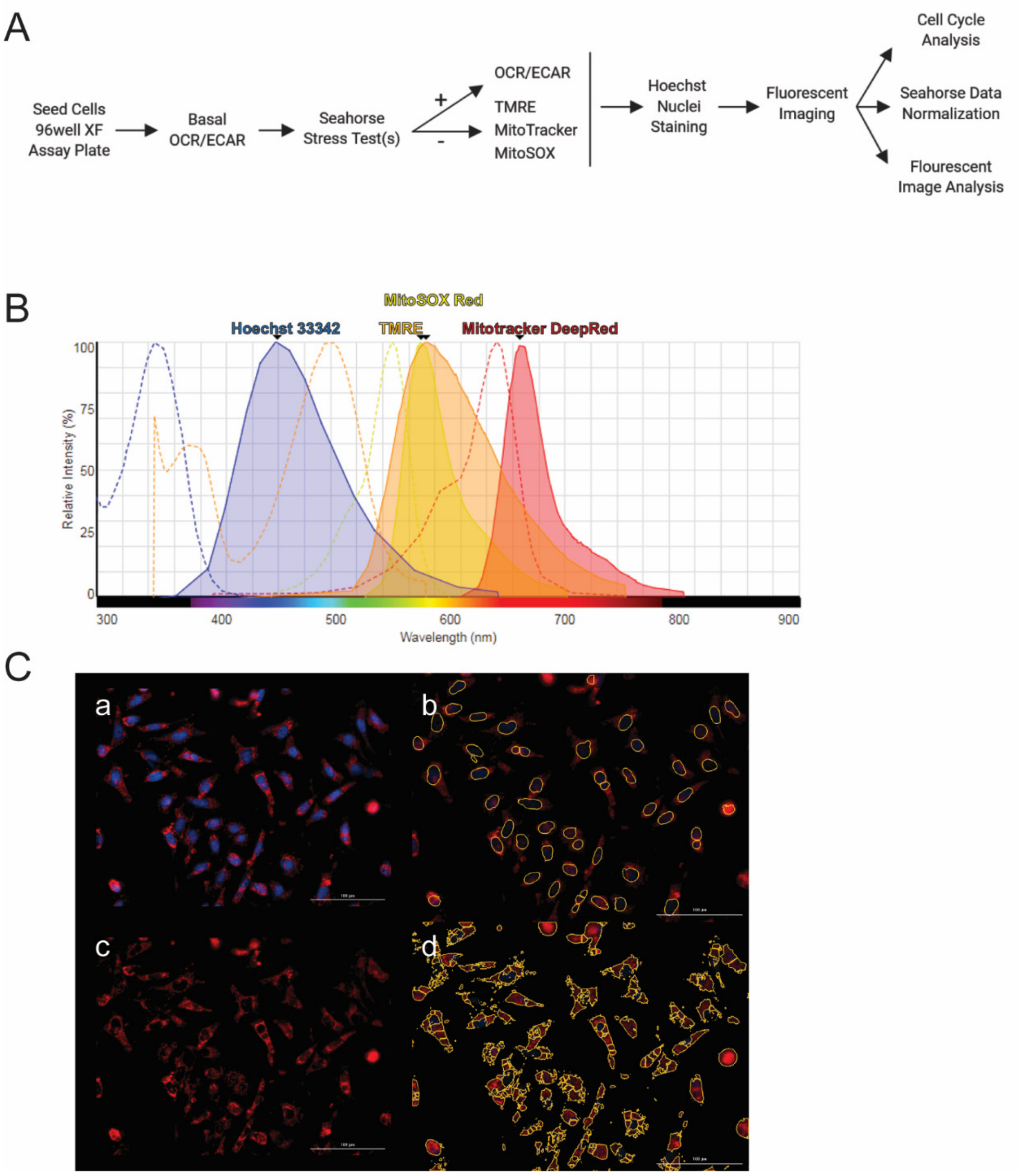


**Supplemental Fig. 1** ***Seahorse-Cytation assay workflow and imaging acquisition and analysis parameters.*** (**A**) Seahorse-*Cytation 5* workflow schematic. (**B**) Emission spectra of the fluorescent stains utilized in this study. (**C**) Hoechst nuclei staining and nuclei masking (a,b) and Mitotracker Red fluorescent staining and its applied secondary mask for image analysis and quantification (c,d).

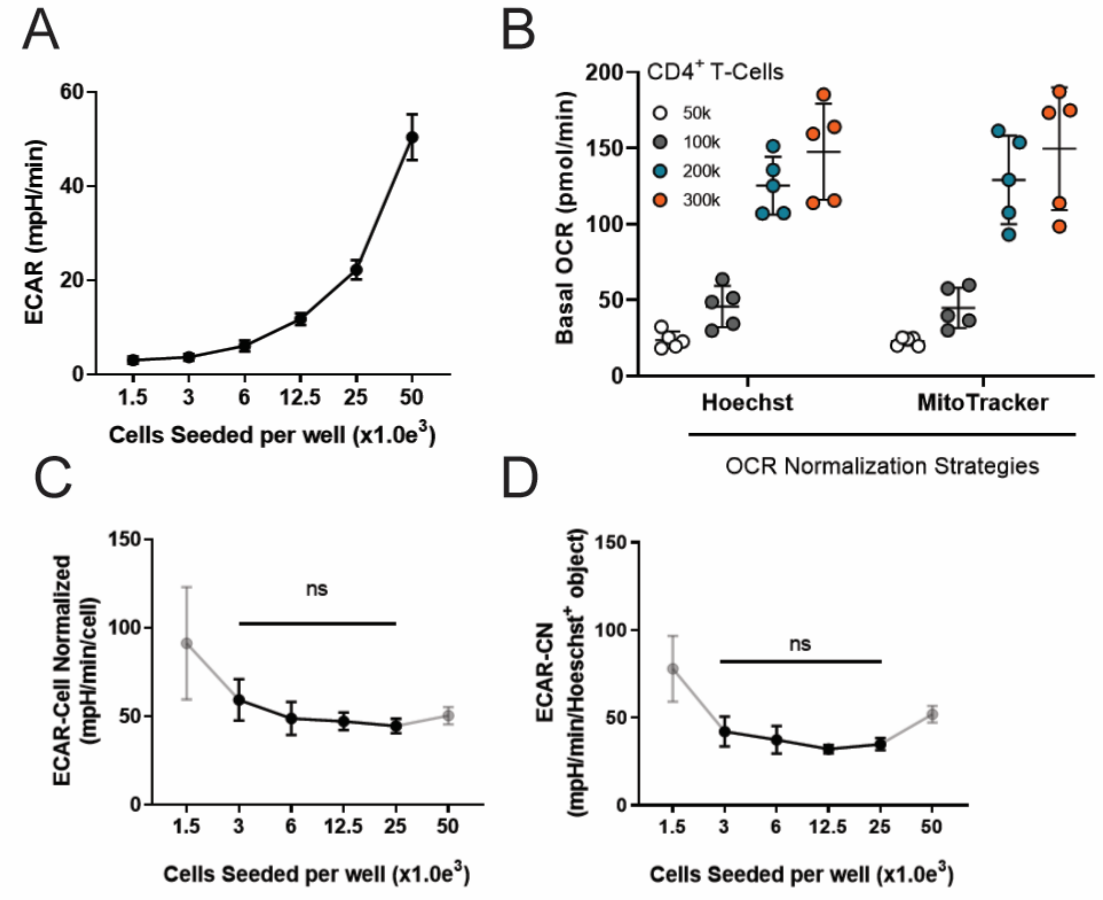


**Supplemental Fig. 2** ***ECAR results based on differing normalization strategies.*** (**A**) Raw ECAR values increase with cell density. (**B**) $CD4^+$ T-cell basal OCR data normalization strategies (i.e. Hoechst – total fluorescently labeled nuclei; MitoTracker – normalization based on mitochondrial units per cell) based on increasing seeded cell densities i.e. 50,000 = 50k, etc. Correcting for cell density with either traditional hematocytometer (**C**) or fluorescent-based nuclei imaging (**D**) has little impact on ECAR data normalization post Mito Stress Test.

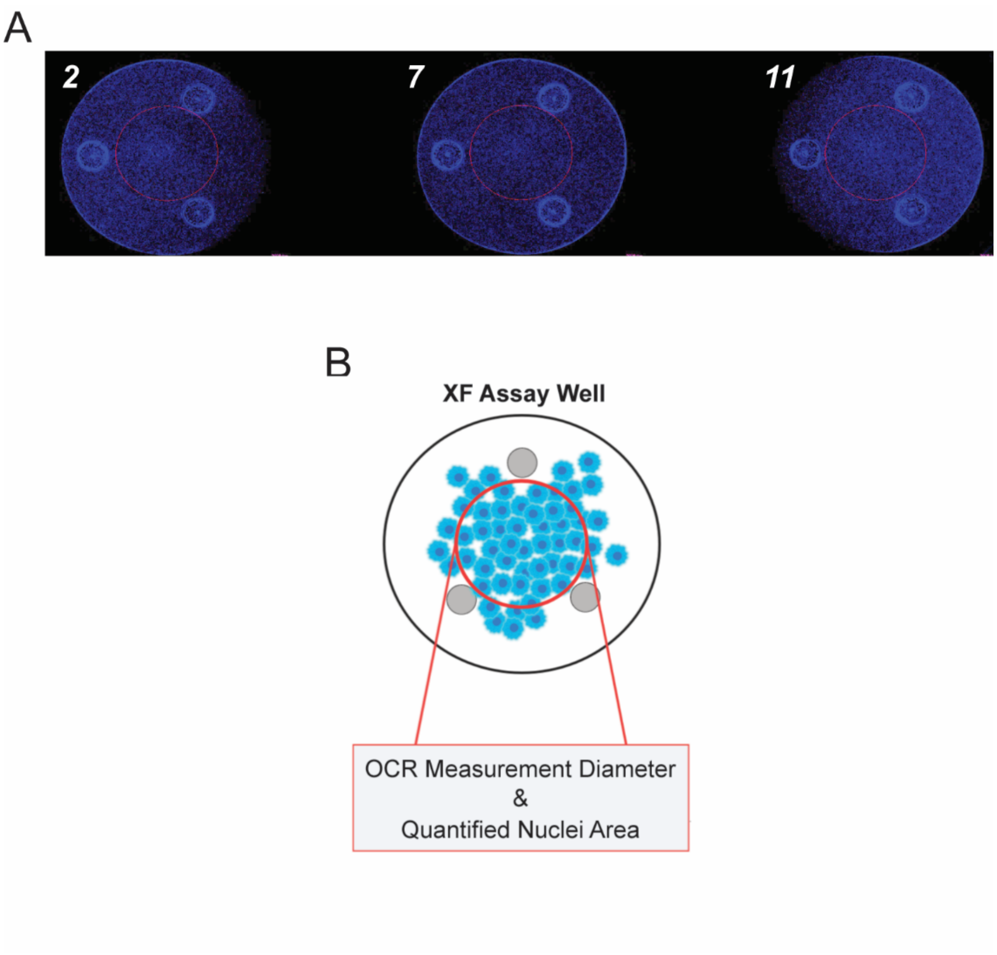


**Supplemental Fig. 3** ***XF assay edge effects and imaging parameters***. (**A**) Representative images of fluorescent nuclei imaging (blue) revealing edge effects in the XF 96 well cell culture plate. Red circle implicates where bioanalyzer data is traditionally captured. Numbers indicate column on XF 96 well plate. (**B**) Schematic representation of a XF 96 cell culture well where respiration measurements and fluorescent nuclei images are obtained. Cells are in blue; sensors are in gray.

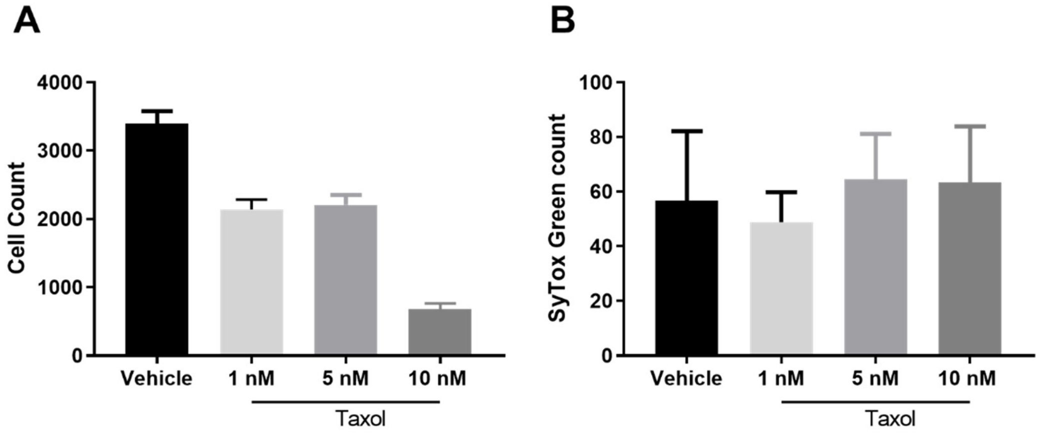


**Supplemental Fig. 4** ***Low dose taxol treatment arrests cell cycle but does not induce significant toxicity***. (**A**) Cell number in taxol treated samples. (**B**) Cell viability by SyTox green staining in taxol treated samples.

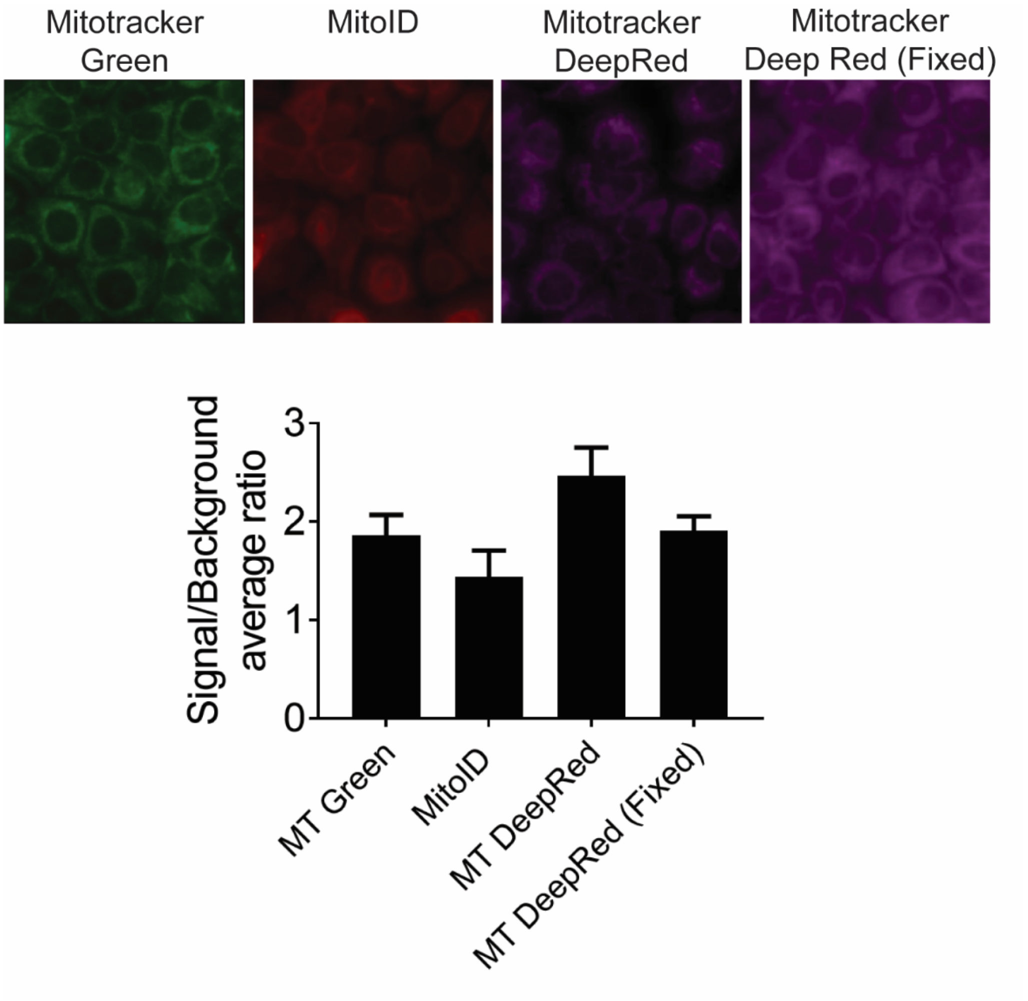


**Supplemental Fig. 5** ***Evaluation of various Mitotracker dyes and determination of their signal to noise background.*** Mitotracker Green, MitoID, and Mitotracker DeepRed were evaluated in living cells while Mitotracker DeepRed was also evaluated post fixation (i.e. “Fixed”). Representative fluorescent images at top. The quantified signal to background ratio is presented at bottom.

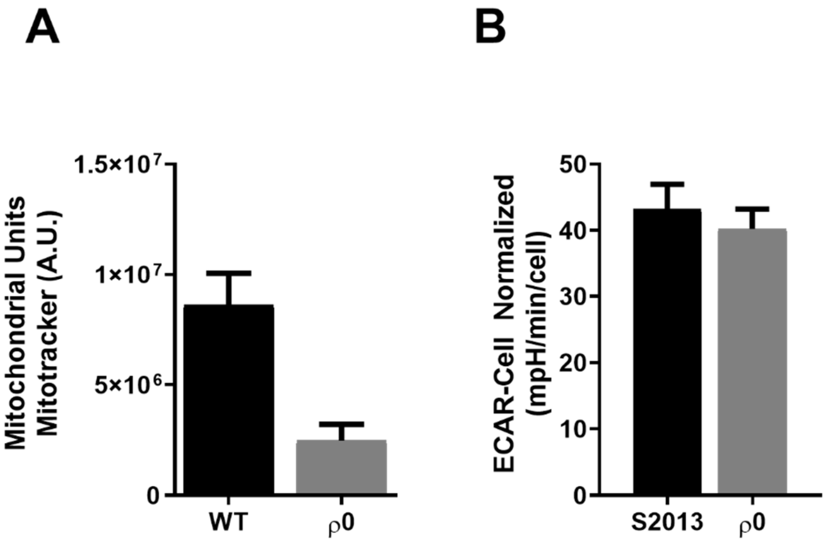


**Supplemental Fig. 6** ***Rho0 S2-013 cells (S2-013-ρ0) mitochondrial quantitation and ECAR values***. (**A**) S2-013 wild type (WT) pancreatic cancer cells harbor greater numbers of mitochondria than their S2-013-ρ0 (Rho0) counterparts, as quantified by total Mitotracker fluorescence intensity. Data displayed as arbitrary fluorescent units (A.U.). (**B**) ECAR measurements in S2-013-ρ0 vs WT cells. Data corrected for overall cell numbers.

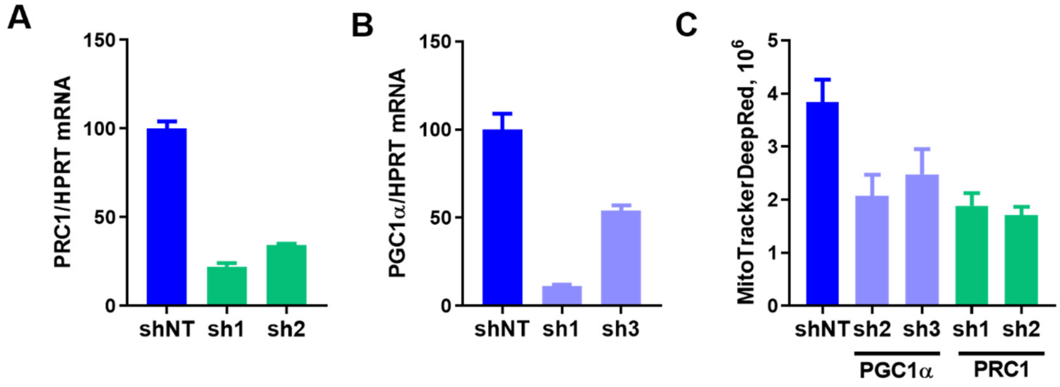


**Supplemental Fig. 7** ***PRC1 and PGC-1a RNAi knockdown characterization.*** Confirmation of PRC1 (**A**) and PGC-1a (**B**) knockdown by short hairpin (sh)RNA as measured by qPCR in T3M4 shPRC1 and shPGC-1a cells. NT, non-targeting. (**C**) Total levels of mitochondria are lower in PGC1a and PRC1 KO cells as determined by Mitotracker fluorescent staining.

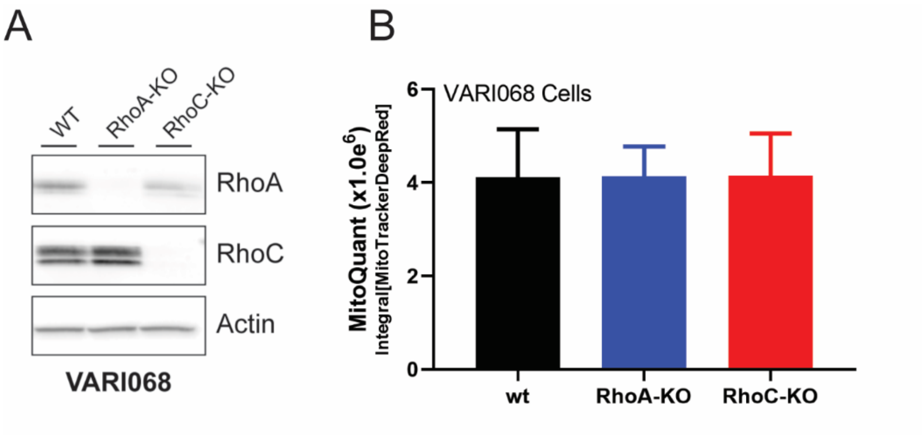


**Supplemental Fig. 8** ***Protein expression and mitochondrial quantification in the Rho knockout VARI068 breast cancer cell model.*** (**A**) Protein expression of RhoA or RhoC in CRISPR knockout breast cancer PDX cell line, VARI068. (**B**) Mitochondrial quantitation as determined by Mitotracker fluorescent staining and imaging in wild type, RhoA KO, and RhoC KO VARI068 cells.

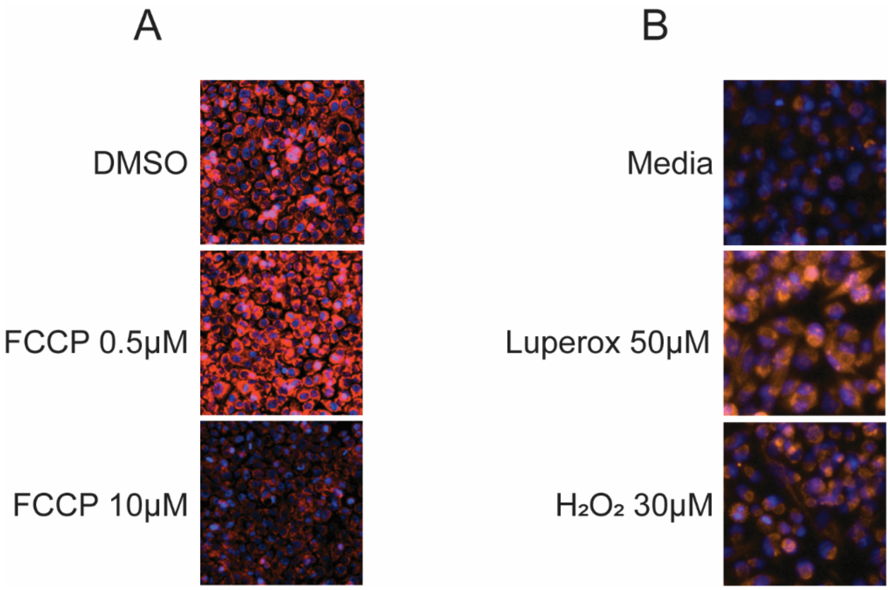


**Supplemental Fig. 9** ***TMRE and MitoSOX fluorescent images***. Representative fluorescent images utilized for image analysis of (**A**) TMRE post FCCP treatment (**B**) or MitoSOX imaging post Luperox/$H_2O_2$ stimulation.

***Supplementary Table 1.*** **shRNA constructs utilized for this study.**

| Denomination | Gene Name | TRCN# |
|---|---|---|
| **PGC1α-sh1** | PPARGC1a | TRCN0000364084 |
| **PGC1α-sh2** | PPARGC1a | TRCN0000364087 |
| **PGC1α-sh3** | PPARGC1a | TRCN0000364086 |
| **PRC1-sh1** | PPRC1 | TRCN0000063296 |
| **PRC1-sh2** | PPRC1 | TRCN0000437395 |
| **NT** | Non-Targeting | CCG GCA ACA AGA TGA AGA GCA CCA ACT CGA GTT GGT GCT CTT CAT CTT GTT GTT TTT |

*Supplementary Table 2.* **RT-qPCR primer sequences utilized for this study.**

| | |
|---|---|
| **HPRT-Fw** | TGACACTGGCAAAACAATGCA |
| **HPRT-Rv** | GGTCCTTTTCACCAGCAAGCT |
| **PARGC1A-Fw** | ACCATATTCCAGGTCAAGATCAA |
| **PPARGC1A-Rv** | GGCTTGACTCATAGTAATAGCAGGA |
| **PPRC1-Fw** | TGGCATCACTTGAGGATGAG |
| **PPRC1-Rv** | AGCCCTCAGGCAGTGTCA |
